# Habitat quality, not patch isolation, drives distribution and abundance of two light-demanding butterflies in fragmented coppice landscapes

**DOI:** 10.1101/2022.12.19.520996

**Authors:** Anne Graser, Marit Kelling, Rebecca Pabst, Meike Schulz, Johannes Kamp

## Abstract

Coppice forests are socio-ecological systems especially rich in biodiversity. They have been transformed into high forest and abandoned across large areas of Europe over the past 200 years. Coppice loss is likely an important driver of insect declines. It is currently unclear whether habitat quality or decreasing connectivity of the remaining fragments is more important for the survival of insect populations. We related the abundance of two coppice-attached butterflies of conservation concern, *Satyrium ilicis* and *Melitaea athalia*, to indicators of habitat quality and habitat connectivity. We estimated butterfly densities using Distance Sampling along a successional gradient (time since last cut: 1–9 years; N = 130 plots) across one of the largest remaining simple oak-birch coppice landscapes in Central Europe. Both species reached abundance peaks within four to six years after the last cut, declining rapidly in abundance with subsequent succession. We found no evidence that coupe size, coppice availability and patch (= coupe) connectivity were related to the density of the species. Besides stand age, the cover of larval foodplants explained predicted butterfly densities well. Only *Satyrium ilicis* benefitted from high Red Deer densities.

Implications for insect conservation: Our results suggest that habitat quality and sufficient availability of coppice of suitable age matters more than coupe size and fragmentation within a traditional managed coppice landscape. Coppice restoration aiming at the study species should ensure a shifting mosaic of successional habitat to provide a large availability of resprouting oak stools and blueberry vegetation that holds dense *Melampyrum pratense* stands.

## Introduction

Habitat loss through anthropogenic land-use change is the most important driver of biodiversity decline globally (Jaureguiberry et al. 2022). Habitat loss has large, consistently negative effects on biodiversity (Fahrig 2003). The effects of habitat fragmentation are less clear. Habitat fragmentation is often defined as the division of habitat into smaller and more isolated fragments, separated by a matrix of human-transformed land cover (Haddad et al. 2015). In a global meta-analysis of experimental studies, habitat fragmentation has been shown to reduce biodiversity by 13 to 75% (Haddad et al. 2015). But habitat fragmentation, once controlled for the loss of habitat amount (habitat fragmentation ‘per se’), can also result in biodiversity gains and positive population responses (Fahrig 2017). However, definitions and approaches to study habitat fragmentation vary across studies, and are therefore difficult to interpret and compare (Fischer and Lindenmayer 2007). There is considerable debate about the effects of habitat fragmentation on biodiversity (Fahrig 2017; Fletcher et al. 2018; Fahrig et al. 2019).

In applied conservation, e.g. the management of Protected Areas, it is often of interest to estimate effects of habitat fragmentation versus those of the loss of habitat amount, or habitat quality. Understanding whether total habitat amount, patch size or habitat quality contribute most to species richness and abundance allows conservation managers to develop appropriate strategies, assessing trade-offs between improving habitat quality and fragmentation of habitats. Habitat quality has repeatedly been shown to be a more important driver of abundance and occurrence than fragmentation, i.e. habitat isolation (Poniatowski et al. 2018; Silva et al. 2022).

Habitat loss and fragmentation are of especially high relevance in forested landscapes (Haddad et al. 2015). Across Central Europe, forest and woodland management has changed strongly over the past 200 years (Kirby and Watkins 2015). With the arrival of coal, oil and gas, the demand for firewood both for industrial and domestic use has been declining since 1800 (McGrath et al. 2015). In the same period, forest use intensity declined: Coppice forests with short rotation cycles were converted into high forest at a large scale (Kirby and Watkins 2015; Kamp 2022), wood pastures were abandoned and afforested, and livestock grazing disappeared from forests in most regions (Plieninger et al. 2015). During the past 50 years, “close-to-nature” silviculture meant an abandonment of clear-cutting practices, additionally resulting in more continuous forest cover over longer periods (Palmero-Iniesta et al. 2021).

The decrease in biomass harvest from forests over the past 200 years (Schelhaas et al. 2003; McGrath et al. 2015) has had important, yet only partly quantified consequences for biodiversity. Historical, open coppice and grazed forests promoted the occurrence of many warm-adapted species that prefer light conditions (Vacik et al. 2009; Weiss et al. 2021). With an increase in biomass, tree-cover and tree-height, and an associated cooler forest microclimate and denser herb layer, many of these species have been declining dramatically over the past century (Hodgson et al. 2009; Swanson et al. 2011; Hilmers et al. 2018). Recently well-publicized declines in birds (Gregory et al. 2019; Burns et al. 2021; Kamp et al. 2021) and insects (e.g. Seibold et al. 2019) over the past decades are at least partly attributable to the abovementioned changes in forest management (e.g. Thorn et al. 2015; Roth et al. 2021; Laussmann et al. 2021; Habel et al. 2022). However, forest management changes as drivers of long-term insect declines have rarely been discussed (e.g. Wagner 2020).

A forest management type that was especially affected by large area losses and associated fragmentation is coppice, both with and without (henceforth “simple coppice”) standards (Buckley 2020; Slach et al. 2021; Kamp 2022). Afforestation of coppice with fast-growing conifers or European beech, but also coppice abandonment, resulted in fast canopy closing and cooler, darker stand conditions. Along with habitat loss, fragmentation of coppice was widespread in the process of transformation to high forest (Deconchat and Balent 2001), as the remotest and least productive areas were transformed or abandoned, and as ownership structures are often complex (Kamp 2022).

Despite these dramatic reductions in coppice area, little is known about biodiversity responses to this century-long coppice loss. Traditionally coppiced woodlands create a small-scale mosaic of different successional stages and therefore provide habitats for different species and their individual requirements (Hilmers et al. 2018). Compelling evidence for community changes and abundance declines due to coppice loss and abandonment is available for plants (Kopecký et al. 2013) and birds (especially long-distance migrants, Fuller and Rothery 2013). Therefore, coppice loss is considered a major driver of the decline of woodland butterflies in Europe (Warren and Key 1991; Bergman 2001; van Swaay et al. 2006). Remaining coppice is considered as a highly valuable habitat especially for butterflies and moths (Fartmann et al. 2013; Roth et al. 2021) as well as saproxylic beetles (Weiss et al. 2021). Coppice restoration and targeted management can lead to biodiversity gains, e.g. in spiders (Vymazalová et al. 2021), moths and butterflies (Hodgson et al. 2009; Dolek et al. 2018a; Roth et al. 2021).

Much less is known about the effects of fragmentation, i.e. increasing isolation of remaining, used coupes on biodiversity, and on the relative contributions of habitat quality on occurrence and abundance of animals and plants in coppice. Interactions with important components of coppice systems, e.g. high Red Deer *Cervus elaphus* densities (Feber 2001; Spitzer et al. 2008; Ramirez et al. 2019), have also rarely been considered.

We used two butterfly species, *Satyrium ilicis* (Ilex Hairstreak) and *Melitaea athalia* (Heath Fritillary), that prefer coppice forests (Warren 1991; Hermann and Steiner 2000; Hodgson et al. 2009; Maes et al. 2014), as model species. We aimed to evaluate whether their populations are currently driven by patch size and fragmentation of remaining coppice patches (= coupes), or by habitat quality and successional stage, as other studies suggest (Warren 1987a, b; Brereton 2006; Ulrich and Caspari 2007; Bräu et al. 2013; Maes et al. 2014). Our main objective was to provide evidence for coppice managers and conservationists about the most important factors to consider when maintaining, managing and restoring coppice. We surveyed these species in what is perhaps the last intact, large-scale remaining simple coppice landscape in Central Europe (Kamp 2022).

We hypothesized that butterfly densities:

1. increase with increasing patch (= coupe) size and increasing patch connectivity, because larger patches exhibit a higher resource availability, and isolation impairs colonization of empty patches.
2. increase with warmer microclimate, because coppice insects are warm-adapted species.
3. peak at species-specific successional stages, because these provide a maximum of resources for larval and adult stages.
4. increase with an increasing habitat quality, i.e. an increasing cover of larval food and nectar plants.
5. increase with increasing intensity of Red Deer grazing, because intensive grazing delays succession towards closed canopy.

## Material and Methods

### Study area

The study region is located in the federal states of Hesse and North Rhine-Westphalia (NRW) in Germany, centred on 50°47’ N, 8°14’ E and 50°58’ N, 8°07’ E. It forms the largest remaining areas of simple coppice (coppice-without-standards) in Germany and is one of the largest in Central Europe. The total size of the study area is 14000 ha (situated between 300 and 600 m a.s.l.), with about 5000 ha active coppice (Kamp 2022). In Hesse, coupes are larger and rotation cycles of recurring cutting are shorter than in North Rhine-Westphalia (Kamp 2022). The parcels are mainly located on slopes and the coppices grow on soil acidic substrates (podzols and brown earths) (Stegger and Vinnemann 2013). Alongside coppice, managed high beech forests and Norway spruce plantations are found in the study area. The climate is characterised by an annual precipitation of 750–950 mm and a mean annual temperature of 8.9 to 9.7 °C (Deutscher Wetterdienst (DWD) 2017).

Coupes are sparsely vegetated in the first two years after cutting. A shrubby stage persists from the third to the ninth year that is dominated by resprouting birch, scotch broom and heather. Approximately 15 years after cutting, the canopy closes to stands dominated by oak and birch, with blueberry carpets in the shrub layer (Suppl. Material Fig. S1). The study area is also characterised by intensive game management and very high densities of Red Deer and Wild Boar, which are typical of coppice forests (Joys et al. 2004; Benes et al. 2006).

### Butterfly surveys

Information on coupe size and the year of the last cut was taken from an area-wide database that was assembled from high-resolution remote sensing data (Kamp 2022). In the study area, coupes that were cut 1–9 years ago were considered as suitable habitat, since early and mid-successional woodland stages are primarily colonized by *S. ilicis* and *M. athalia* (Fartmann et al. 2013; Maes et al. 2014). In 2018, 86 coppice woodland patches were selected randomly in Hesse, stratified by size, isolation and stand age of the coupes. In the following year, we revisited 38 out of the 86 patches and added 44 new patches, 4 fresh cuts in Hesse (1 year after cutting) and 40 plots in NRW (1–9 years after cutting) (Fig. 1). The single coupes varied in size between 0.32 and 17.77 ha (mean ± SD: 5.10 ± 4.18) in Hesse and between 0.38 and 5.42 ha (mean ± SD: 1.73 ± 1.17) in NRW. At each coupe, a starting point was chosen randomly (with a buffer of 50 m to the edge to avoid strong edge effects), and a transect was walked set from this starting point dissecting the coupe in the direction of the slope. Due to varying coupe size, transects varied in length from 40 to 455 m (mean ± SD: 161.82 ± 94.21).

**Fig. 1.**
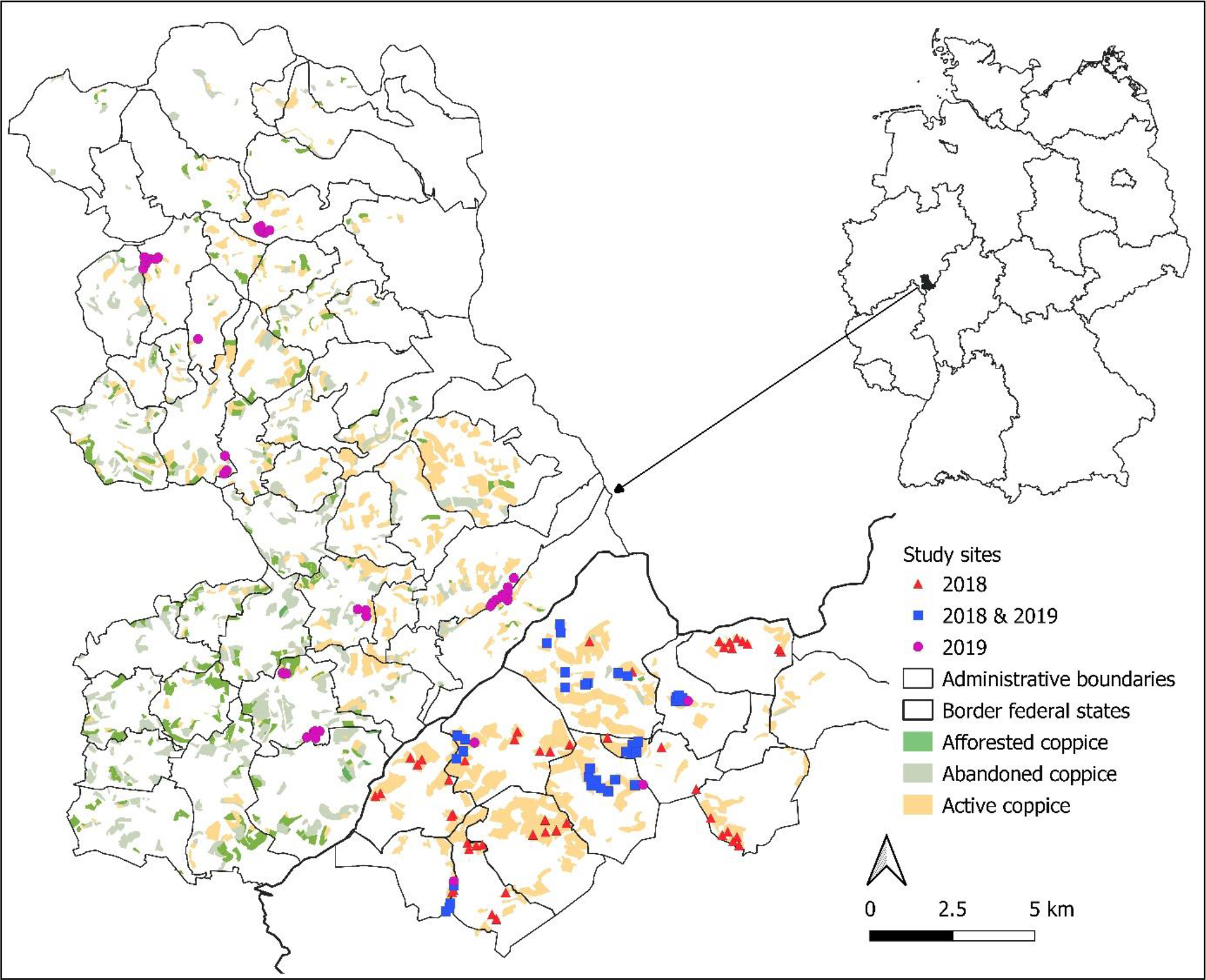
Location of the study area in Germany with all remaining active coppice coupes (yellow), afforested coppice (dark green) and abandoned coppice (light green) (Kamp 2022). Investigated coupes are marked with different symbols

We surveyed all transects for *S. ilicis* and *M. athalia* five times per year between late May and mid-July in 2018 and 2019. To correct for varying factors that affect detectability, and to predict population densities without labour-intensive capture-recapture efforts, we used Distance Sampling rather than simple Pollard walks (Buckland et al. 2008; Isaac et al. 2011). Observations were assigned to five distance classes (0–2.5 m, 2.5–5 m, 5–10 m, 10–20 m, >20 m) on both sides of the transect line. We identified *S. ilicis* and *M. athalia* with the aid of binoculars. Occasionally, hand-held nets were used to catch individuals and release them afterwards. Surveys took place between 9:00 a.m. and 7:00 p.m. (GTM + 2), at wind speeds below 4 Bft. and temperatures >17 °C when the sky was clear, at least 20 °C at less than 40 % cloud cover, and at least 25 °C when overcast.

### Habitat quality

To assess habitat quality for *S. ilicis* and *M. athalia* we collected information on parameters known to influence their occurrence and abundance. In late June of each survey year, parameters were recorded at the beginning, midpoint and end of each transect, in a quadrat of 10 × 10 m. Endpoints were placed 10 m before the end of the transect to avoid strong edge effects. For analysis, measurements were averaged over the three recording points. We measured cover and heights of herb, shrub and tree layer for each coupe. To assess the availability of preferred host plants for the larvae and nectar source for butterflies, we estimated the cover of *Quercus spp*. (both *Quercus robur* and *Quercus petraea*) in the area, which is the foodplant of *S. ilicis*, *Melampyrum pratense* and *Digitalis purpurea*, which are larval foodplants of *M. athalia*, and *Rubus fruticosus* agg. that is a key nectar resource for both species in the area. We estimated relative Red Deer grazing intensity by counting dung piles at a strip of 2 m width along the transects, and later calculated dung density per 100 m^2^ (Marques et al. 2001).

For each plot, terrain height, slope and aspect were calculated from a digital elevation model (U.S./Japan ASTER Science Team 2011) in QGIS Version 3.14 and then used to calculate a heat load index following McCune (2007).

### Habitat fragmentation and landscape matrix

To evaluate the fragmentation of coppice on a landscape scale the area of each sampled coupe and that of neighbouring coupes suitable for colonization (stand age 1–9 years) within a 2 km radius were determined. Together with the Euclidean distance from the edge of each coupe to all edges of neighbouring coupes, measured by the use of R package “rgeos” (Bivand et al. 2017), we calculated the habitat connectivity (*I*) of each sampled coupe (*i*) by the following formula:

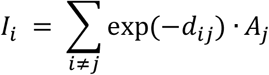

with *Aj* being the size of the neighbouring coupe *j* and *dij* the Euclidian distance from the focal coupe *i* to the neighbouring coupe *j* (Steffan-Dewenter and Tscharntke 2000). We used Euclidian distance instead of functional distance to calculate connectivity because of the lack of species-specific information on resistance values for landscape elements (Poniatowski et al. 2016). A high value of *I* indicates high connectivity and low fragmentation. To characterise the surrounding landscape of each coupe, the percentage cover of high forest, transitional woodland/shrub and grassland/pastures within a 1000 m radius of the coupe was determined (Berg et al. 2011) based on Corine Land Cover 2018 (European Union, Copernicus Land Monitoring Service 2019).

### Data analysis

All statistical analyses were performed in R version 4.1.3 (R Core Team 2022). We first modelled butterfly densities for each sampling site with Hierarchical Distance Sampling models (Royle et al. 2004) using the function *gdistsamp* of the package “unmarked” (Fiske et al. 2015). We used a half-normal detection probability function because we had comparatively few distance classes. We tested variables influencing the detectability of butterflies (e.g. weather, observer) in the detection part of the *gdistsamp* function. As models containing these were consistently not selected as best models, we decided not to include covariates in the detection part of the model. We also estimated abundance independently of covariates, i.e. with an intercept-only model. For subsequent analyses, we used the estimated density of that survey round that yielded the highest density (i.e. represented the phenological peak of the flight season).

Since hierarchical Distance Sampling models do not easily allow for an inclusion of random effects, but as this was necessary due to spatial dependencies in the data, we extracted transect-level predicted densities of both species and related them in a following step to explanatory variables. To predict abundance as a function of habitat parameters and isolation, we used Generalised Linear Mixed-effects Models (GLMM) fitted with functions of the “glmmTMB” package (Magnusson et al. 2017). The response variable, modelled population density per transect, is a continuous variable. We compared the performance and goodness-of-fit of models assuming different distributions (gaussian, gamma, tweedie) using the Aikaike information criterion (AICc) and a visual inspection of the residuals. Models using the tweedie distribution performed consistently better than those assuming other distributions, therefore we fitted the final models with this distribution. We incorporated study sites nested in communal district as a random effect in all models to account for district-level spatial dependencies among samples that are due to pronounced district-level variation in coppice management (Kamp 2022) and variation in topography and geography. Explanatory variables were standardised and also entered as squared variables to allow for hump-shaped relationships. To avoid multicollinearity, only one of a pair of correlated variables (Spearman’s rho >|0.6|) was allowed in the same model. We used the “DHARMa” package to evaluate the models and to test if model assumptions were met (Hartig 2020). To compare the models the function *model.sel* of the package “MuMln” was used (Bartoń 2022). Conditional and marginal R^2^ were calculated following Nakagawa and Schielzeth (2013). For predictions, the function *ggpredict* in the package “ggeffects” (Lüdecke et al. 2022) was applied. Marginal effects (i.e. prediction when keeping all other model variables at their mean) of explanatory variables were visualized. We used GLMMs without covariates and coupes nested in communal district as a random effect (with tweedie-distribution) to predict mean density during flight maximum and total population size per species across the investigated coupes. To avoid spurious results and data dredging, we specified a number of separate full models based on the hypotheses outlined in the introduction. We compared these models using AICC and Akaike weights, as they were all fitted on the same dataset.

A Principal Component Analysis (PCA) was used (function *prcomp* in package “stats”) to visualise the relationship between explanatory habitat variables.

## Results

### Occurrence, density and population size

In 2018, *M. athalia* was detected at 28 and *S. ilicis* at 59 coupes (age 1–9 years, N = 86 transects). *M. athalia* occurred during the first two sampling occasions and *S. ilicis* during the entire sampling period (Fig. 5). In 2019, *M. athalia* was detected at 34 sites and *S. ilicis* at 42 coupes (N = 82 transects), flight periods were later than in 2018 (Table 1, Fig. 5). Estimated densities and total population sizes were higher for *S. ilicis* than for *M. athalia* (Table 1).

**Fig. 2.**
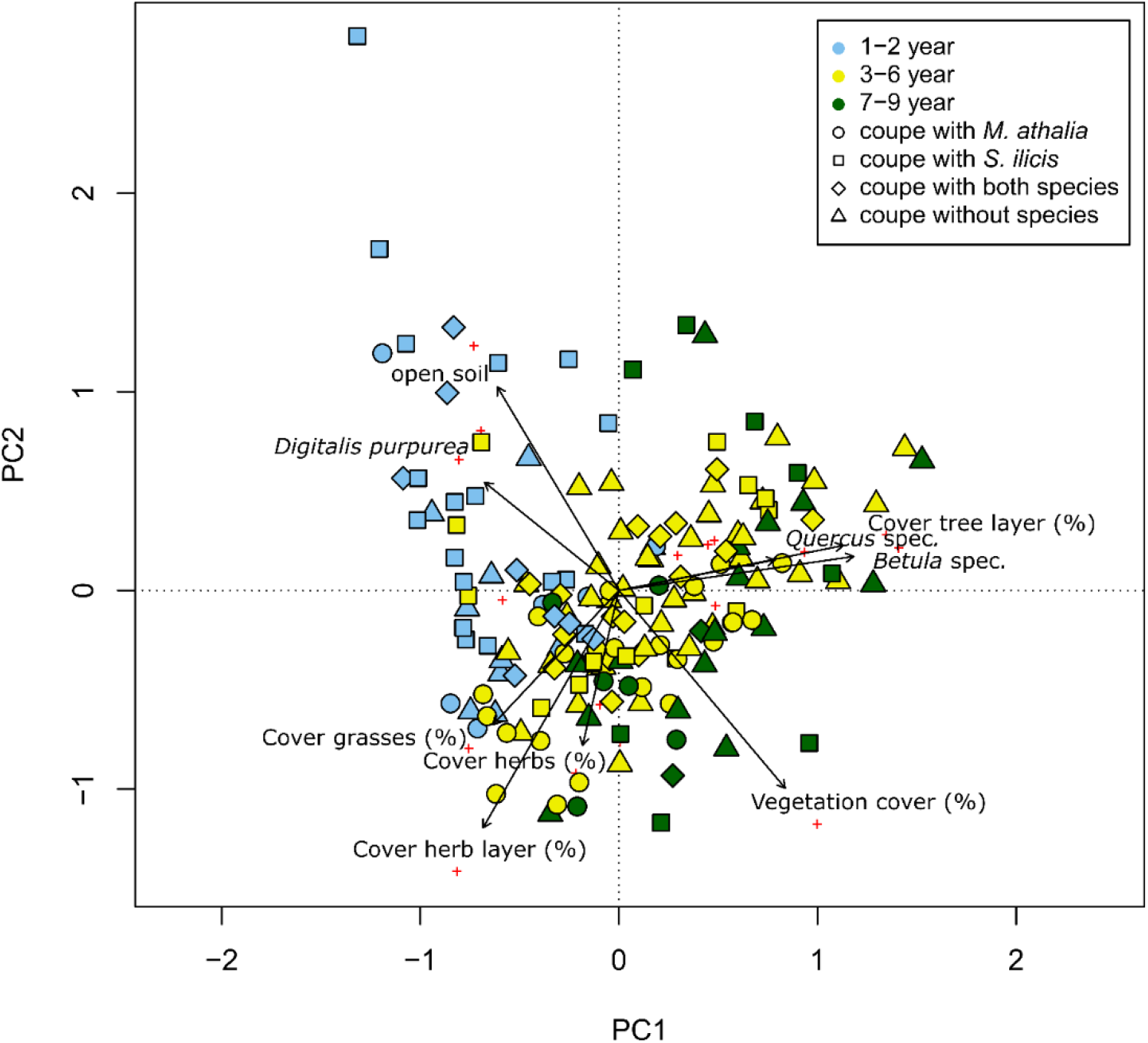
Principal component analysis of habitat parameters. Light blue: early successional stages (1–2 years since last cut); yellow: intermediate stages (3–6 years since last cut); green: late successional stages (7–9 years since last cut). PC1 (Eigenvalue 3.36) and PC2 (Eigenvalue 2.96) together explained 33% of the variance in the data. Habitat parameters with a regression coefficient >|0.8| along the first two axes are depicted. Circles show coupes with *M. athalia* present, squares show coupes with *S. ilicis* present, rotated squares show coupes with both species and triangles show stands with neither species present. Red plus signs show estimated correlation values of each variable with PC1 and PC2 (see also Suppl. Material Table S4)

**Fig. 3.**
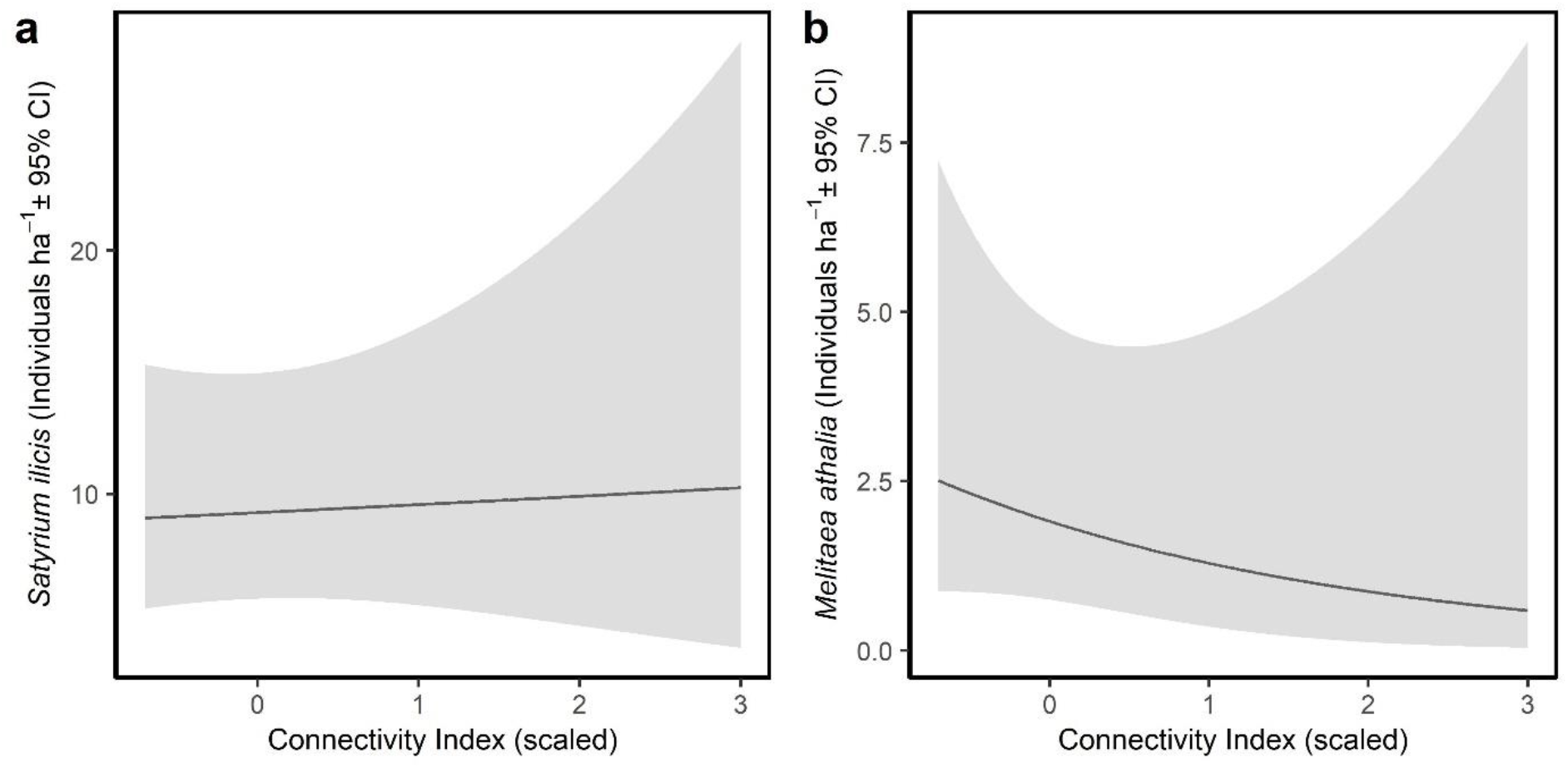
Predicted densities (marginal means ± 95% confidence intervals) of the two butterfly species (a) *S. ilicis* and (b) *M. athalia* as a function of coupe connectivity, from models S.i.1 and M.a.1 in Table 2 (effect not significant). All other covariates of the model were kept constant at their mean

**Fig. 4.**
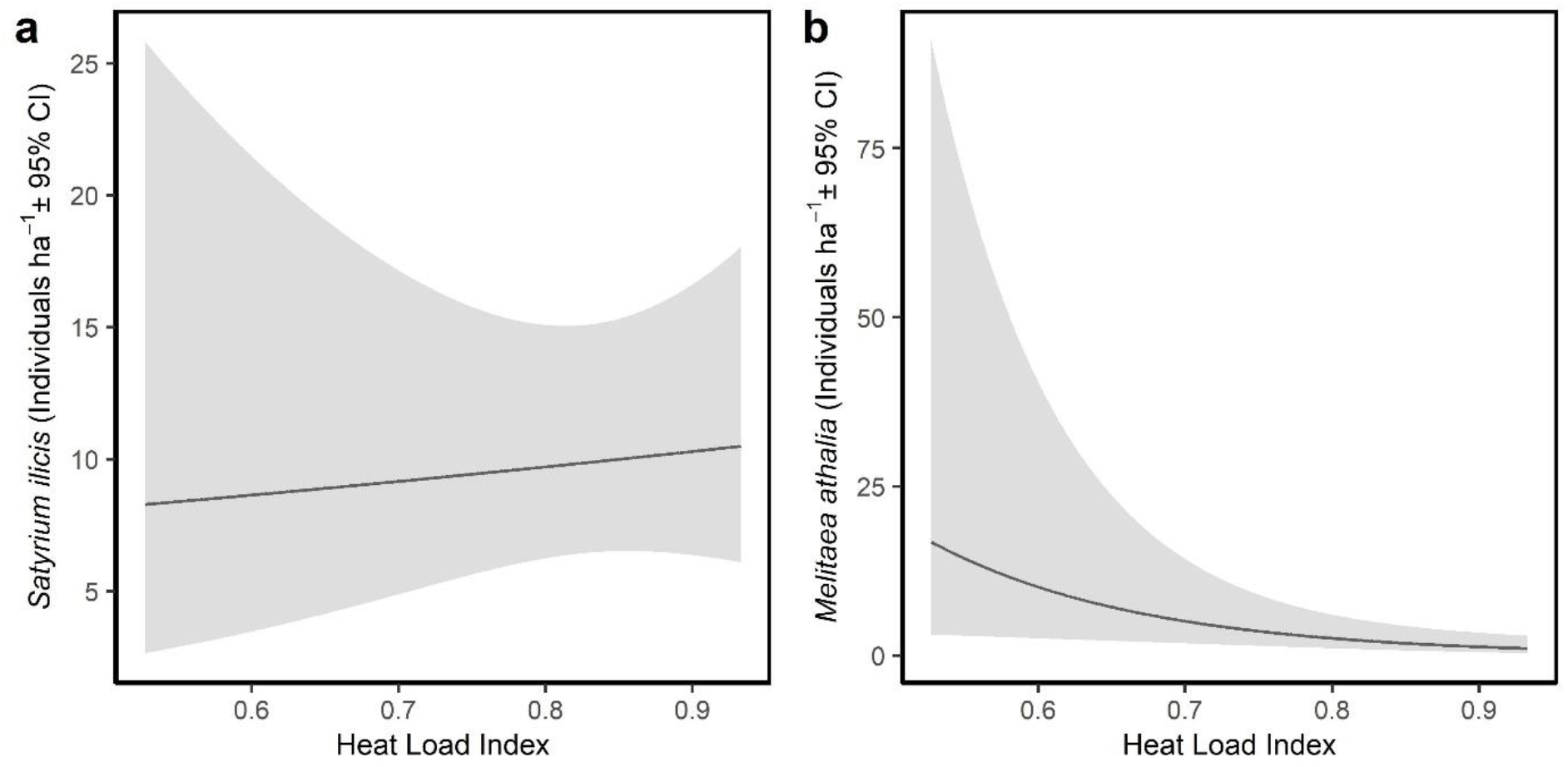
Modelled densities (marginal means ± 95% confidence interval) of the two butterfly species (a) *S. ilicis* and (b) *M. athalia* predicted as a function of the plot-level heat load index, from models S.i.2 and M.a.2 in Table 2 (only significant for *M. athalia*). All other covariates of the model were kept constant at their mean

**Fig. 5.**
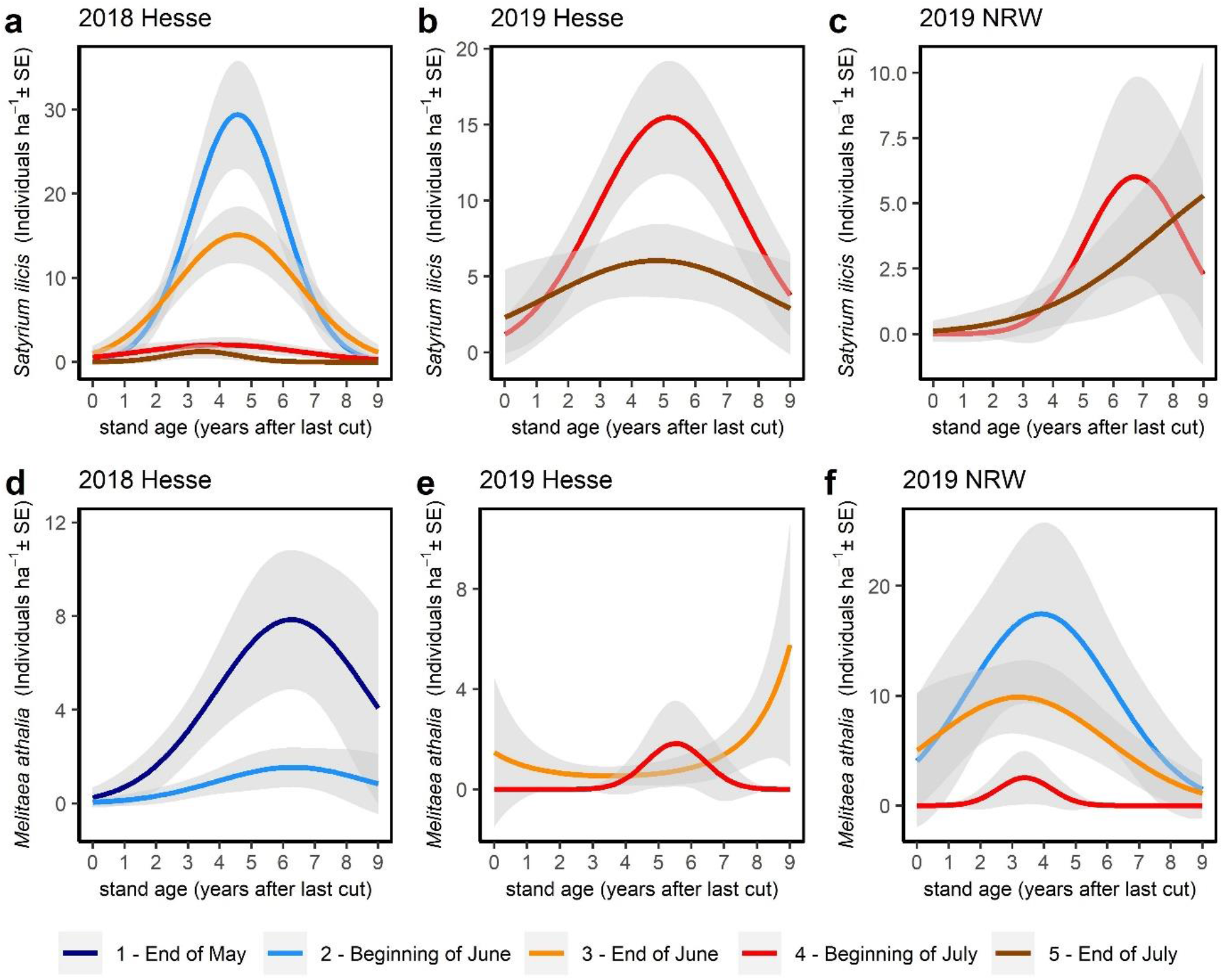
Variation in predicted densities (marginal means ± 95% confidence intervals) as a function of stand age (years since last cut) across all sampled coupes, separately per year and study region, for *S. ilicis* (a, b, c) and *M. athalia* (d, e, f). Phenological variation over the annual sampling periods is also illustrated, with dark blue corresponding to the first survey round, light blue to the second, orange to the third, red to the fourth and brown to the fifth round. All other covariates of the model were kept constant at their mean

**Table 1.** Prevalence, phenology, population densities and total population sizes across the surveyed coupes for *S. ilicis* and *M. athalia* in 2018 and 2019 (*individuals per ha). Total population size estimates are predictions from a GLMM without any covariates. Densities during flight maximum are predictions from Distance Sampling models

| Species | Year | Sample size<br>(number of<br>occupied sites) | Flight period | Density during flight<br>maximum (Mean ± SE)* | Total population size (±<br>SE) during flight<br>maximum |
| --- | --- | --- | --- | --- | --- |
| *Satyrus ilicis* | 2018 | 86 (59) | 28 May –05 July | 16.34 ± 8.36 | 7503 ± 82 |
| | 2019 | 82 (42) | 27 June –22 July | 10.33 ± 7.43 | 3486 ± 66 |
| *Melitaea athalia* | 2018 | 86 (28) | 28 May. –15 June | 3.87 ± 6.87 | 1805 ± 45 |
| | 2019 | 82 (34) | 01 June –03 July | 8.20 ± 15.61 | 2235 ± 63 |

### Environmental conditions across the surveyed coppice coupes

During the first two years after clear-cutting, the coupes were characterized by much open soil, dead wood and a low shrub and tree layer. In mid-successional stages (year 3–6) shrub layer cover peaked, but there was still a distinct grass and herb layer. In the late successional stages (year 7–9), tree cover increased, and the canopy started to close. Fewer shrubs were present (Fig. 2, Suppl. Material Fig. S2, Table S3).

### Habitat models

Neither patch (=coupe) connectivity nor landscape permeability were significant predictors of the density of *M. athalia* and *S. ilicis* (Table 2, Fig. 3). A higher abundance of *M. athalia*, but not *S. ilicis* was associated with a higher heat load (Fig. 4). Both species showed a humped-shaped relationship with time since last cut, for *S. ilicis* the model containing this variable receiving by far the highest support in comparison to all other models (Table 2, Fig. 5). *S. ilicis* increased in abundance with an increase in oak cover and showed highest predicted densities at mean shrub heights between 100 and 130 cm (Fig 6). *M. athalia* densities were positively related to the cover of its larval food plant (*Melampyrum pratense*) (Table 2, Fig. 6). The intensity of Red Deer grazing significantly positively affected *S. ilicis* (Table 2), while *M. athalia* did not show a significant response to grazing (Fig. 7). Model comparisons suggested that for *S. ilicis*, stand age (time since last cut, model S.i.3) explained most of the variation in the data given a certain model complexity based on AICc values (Table 2). For *M. athalia*, in all three models, the stand age model (model M.a.3), the microclimate model (model M.a.2) and the larval food plant model (model M.a.4) had similar fit based on AICc values (ΔAICc ≤2, Table 2).

**Fig. 6.**
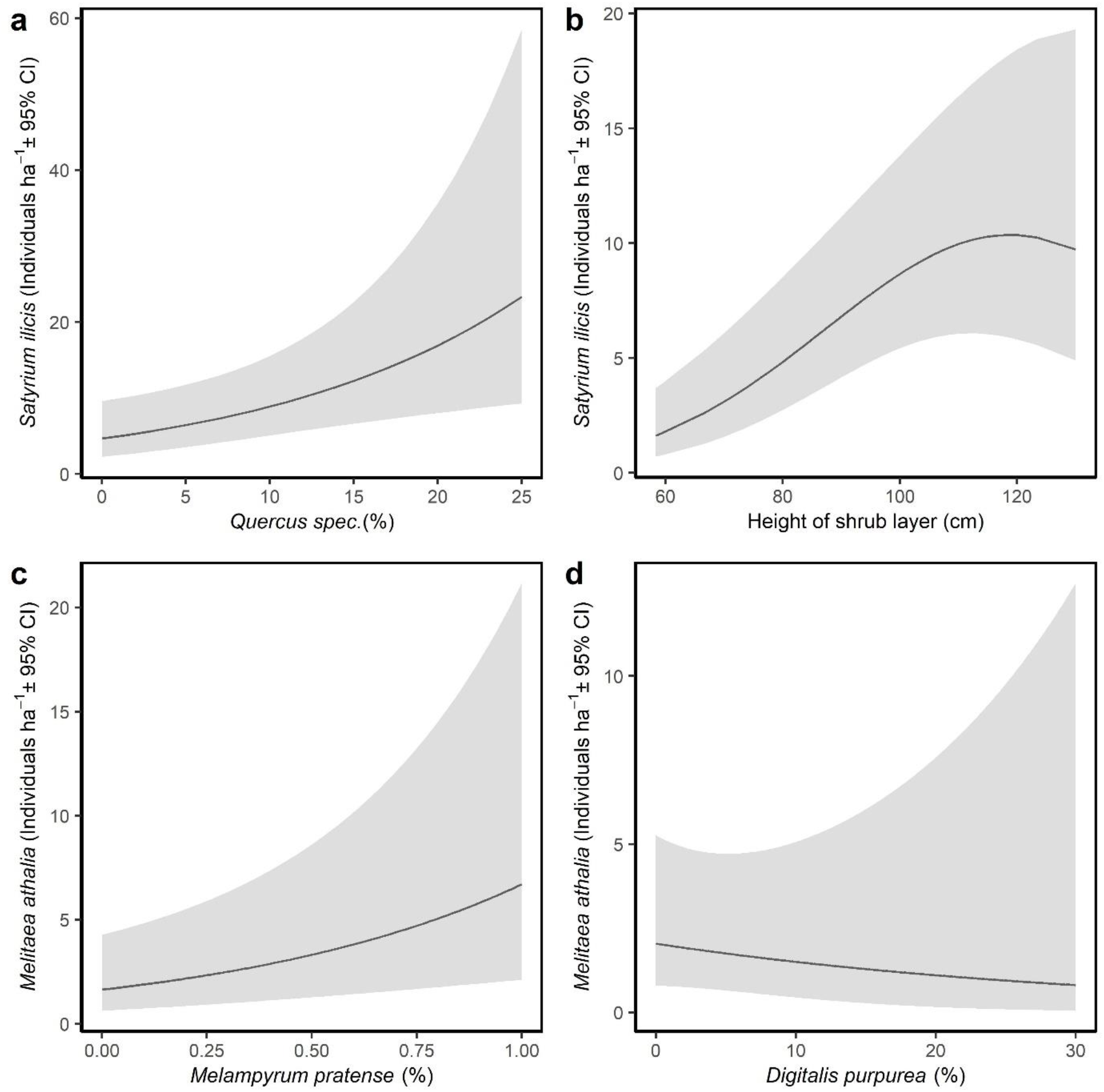
Predicted densities (marginal means ± 95% confidence interval) of *S. ilicis* (a, b) and *M. athalia* (c, d) predicted as a function of the plot-level larval foodplant cover (%), from models S.i.4 and M.a.4 in Table 2. All other covariates of the model were kept constant at their mean

**Fig. 7.**
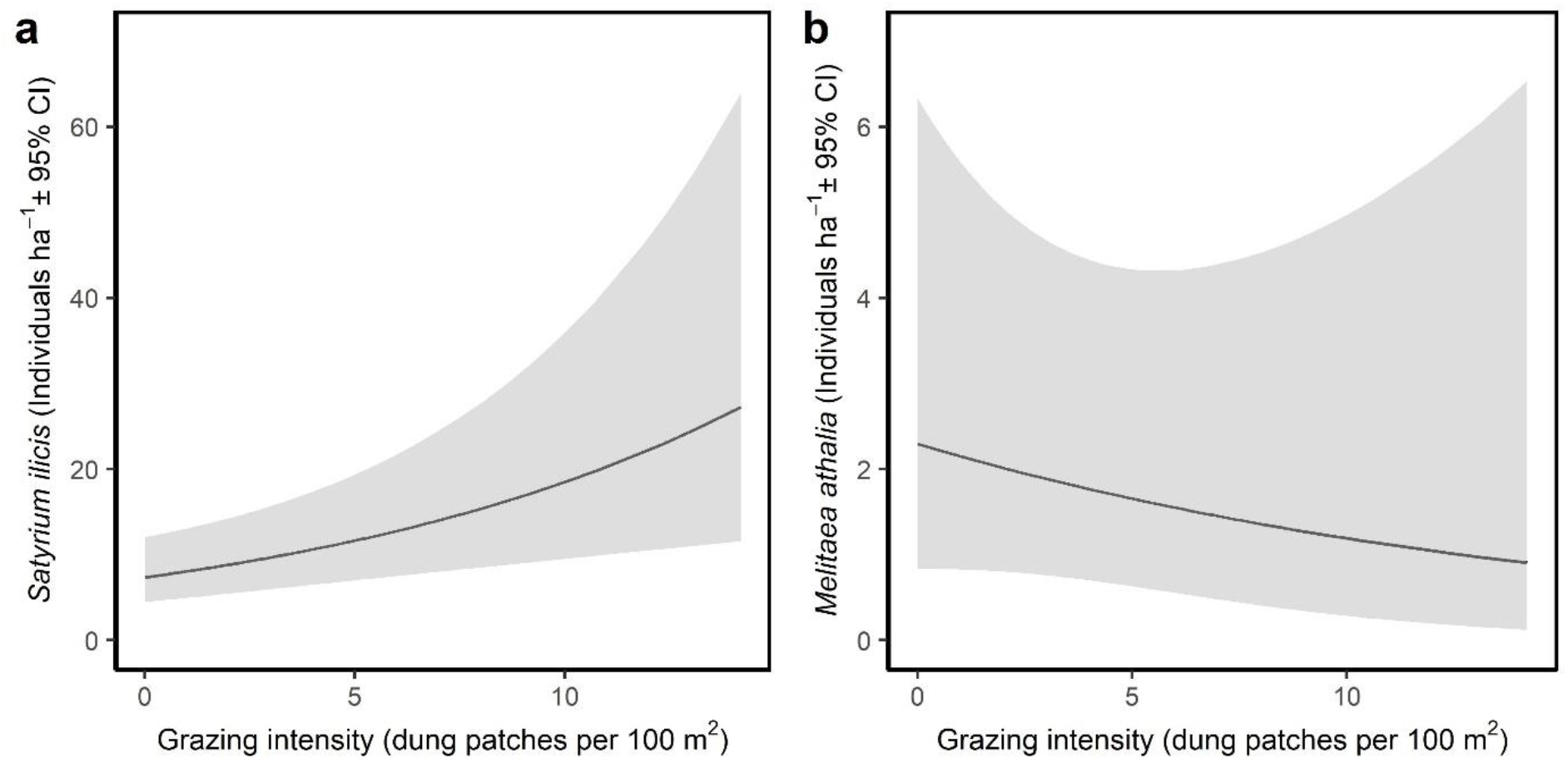
Predicted densities (± 95% confidence interval) of *S. ilicis* (a) and *M. athalia* (b) as a function of Red Deer grazing intensity, from models S.i.5 and M.a.5 in Table 2. All other covariates of the model were kept constant at their mean

**Table 2.**
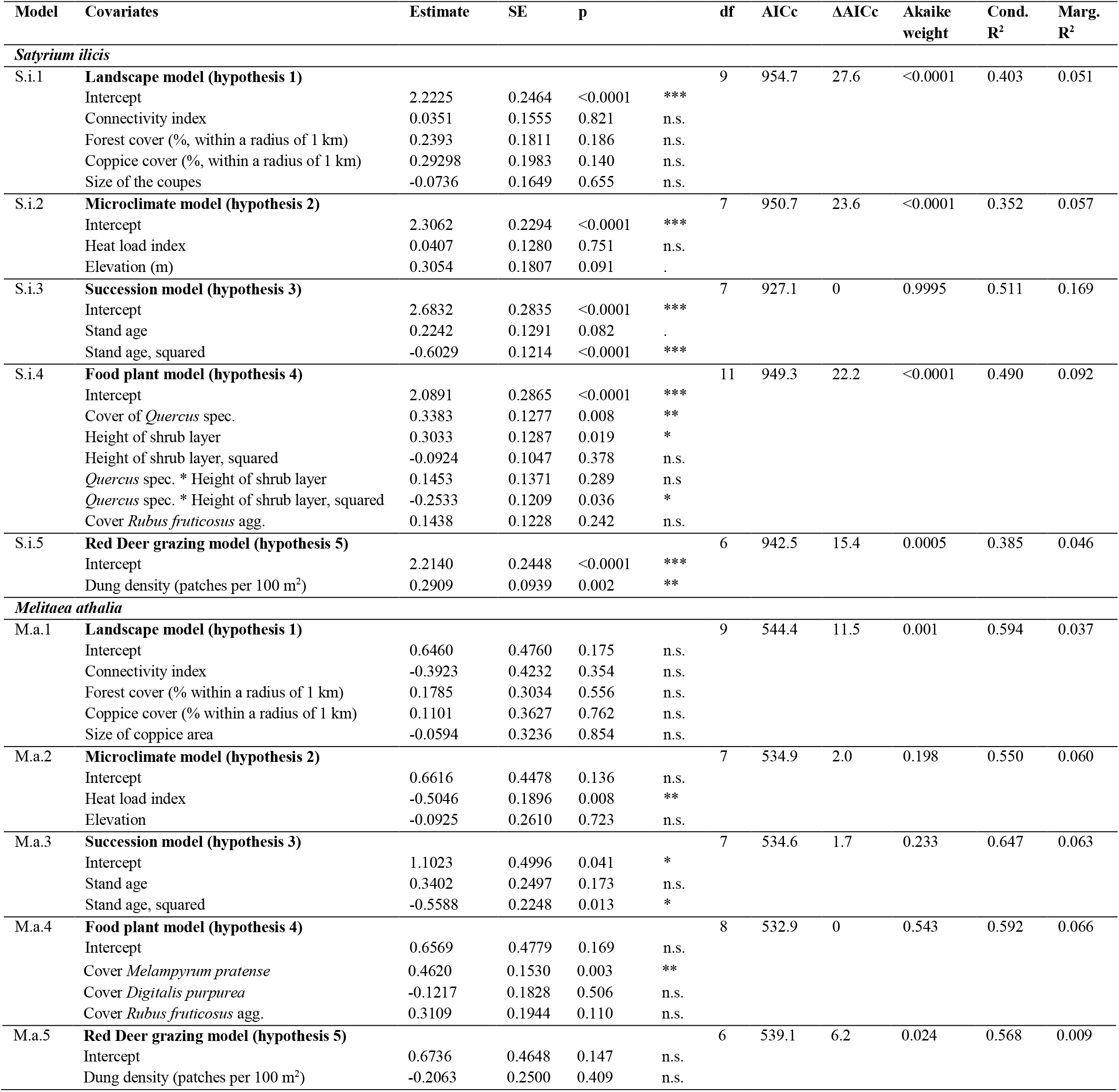
Results of GLMMs explaining modelled densities of the two butterfly species at the sample transect scale. df = degrees of freedom, AIC_C_ = Akaike information criterion adjusted for small sample size, n.s. = not significant

## Discussion

We showed that one of the last intact simple coppice landscapes of Central Europe provides high-quality habitat for *M. athalia* and *S. ilicis* and hosts large populations. The total population size of *S. ilicis* during the flight maximum across the survey coupes was predicted to be 7503 individuals in 2018 and 3486 individuals in 2019. For *M. athalia*, the total number of individuals predicted was 1805 in 2018 and 2235 in 2019 across the surveyed coupes. Occurrence and numbers of individuals were strongly determined by habitat quality and changes therein along a successional gradient, represented by the stand age, i.e. time since last cut. Population density of *M. athalia* imagines increased with increasing cover of the primary larval host plant *Melampyrum pratense* but was negatively affected by a high heat load index. *S. ilicis* population density was positively associated with the intensity of Red Deer grazing, as well as increasing cover and a specific height of oak trees resprouting from stools as host plants for the larvae.

Coupe connectivity showed no measurable effect on modelled densities at flight maximum for either species. Because the availability of suitable habitat in the landscape is still relatively high across our study area, the importance of patch isolation and connectivity might be of secondary importance to the species here (Krauss et al. 2003; Bergman et al. 2004; Krämer et al. 2012). The higher importance of habitat quality over isolation and patch size was also emphasized in other studies (Fleishman et al. 2002; van Halder et al. 2015; Poniatowski et al. 2018) and shown for other butterfly species (Thomas et al. 2011; Krämer et al. 2012; Fartmann et al. 2013).

Climate and weather have a substantial impact on all development stages of butterflies (Radchuk et al. 2013); sometimes even single weather events can affect populations heavily (Lynch et al. 2014; Badik et al. 2015). Consequently, the microclimatic conditions are an important factor influencing habitat quality, which is crucial for the continued survival of metapopulations. Other studies have suggested that *S. ilicis* prefers warm microclimate conditions (Maes et al. 2014). In our study, such a relationship was not detectable. *M. athalia* showed lower densities in areas of a high heat load index. This indicates that the species can make better use of steeper and north-exposed, and therefore cooler slopes. The decreasing density with higher heat stress can be interpreted as an indication of not coping optimally with hot environments, for example early successional stages, and might make the species vulnerable to climate change. About five to six years after the last cut, total cover and denser vegetation create a cooler microclimate, as also Fartmann et al. (2013) found in coppice with standards in French Alsace. In contrast, in south eastern England, where the climate is oceanic, populations of *M. athalia* in coppice forests showed a clear preference for sunny and warm sites (Warren 1987a).

The abundance of both species peaked at a stand age (time since last cut) of ca. four to six years, illustrating the high importance of open coppice during early-succession. Stand age reflects a certain habitat quality, as both vegetation composition, its height and density, and microclimate change with progressing succession. *M. athalia* reached highest densities somewhat later than *S. ilicis*, as also shown in other studies (Treiber 2003). This successional stage is characterised by a rich supply of young oak and birch shrubs, which, however, do not yet dominate and allow for a rich herb layer, where some shaded patches already exist. In later stages of succession after about eight years, habitat quality decreases (Warren and Key 1991; Strätling 2010; Blixt et al. 2015). Other studies also point to peaking butterfly abundances in successional coppice habitats between two and seven years after clear-cut/disturbance (Warren 1987b; Fartmann et al. 2013; Dolek et al. 2018b), suggesting that our results might be generalizable for coppice-dependent butterfly species. Habitat characteristics of stands of the same age, in addition to influences from herbivory (Benes et al. 2006), can vary due to geographic location and therefore show different levels of succession. In comparison to our findings, coppice systems in the UK investigated by Warren (1987b) showed a closed canopy after five to six years and highest abundances of *M. athalia* in stands of age two to four years. In Germany, the study site showed a closing canopy after ten to twelve years, likely because of slower succession processes, and therefore the highest predicted abundances were found in comparatively older stands. The preference of *S. ilicis* for earlier successional stages is evident by its preference of low-growing vegetation and open habitat (Treiber 2003; Strätling 2010; Maes et al. 2014). The traditional, regular management of coppice woodlands in the studied low mountain range of Central Germany is of central importance for *M. athalia* and *S. ilicis*. The annual cutting of areas of varying size creates a heterogeneous mosaic of different succession stages. A sufficient supply of suitable areas nearby is essential for the survival of many light forest/woodland butterflies (Warren and Thomas 1992; Brereton 2006; Twardella and Fasel 2007; Hodgson et al. 2009).

Especially species with monophagous larval stages are strongly bound to their host plants (Krauss et al. 2005; Krämer et al. 2012; van Halder et al. 2015). In general, the occurrence and abundance of many species is particularly influenced by the requirements of the preimaginal stages, as they are little mobile and are mostly relatively long-lived in comparison to the adult butterfly (Fartmann and Hermann 2006; Thomas et al. 2011). For *M. athalia*, a high availability of the larval food plant allowed high densities of the imagines in our study. The availability of *Digitalis purpurea*, as a food plant for the later larval stages, was not a significant predictor of adult population density. Despite the low average cover of *M. pratense*, considerably higher *M. athalia* densities were predicted even at comparatively little increase of cover. *M. pratense* thus acts as a strong limiting factor in our populations, as also shown by Warren et al. (1984). Oak cover was not among the most important parameters in explaining the density of *S. ilicis*. This may be due to the generally high stand densities of oaks in the study area and the huge number of resprouting oak stools (likely hundreds of thousands). The higher densities of *S. ilicis* at shrub heights between 100 and 130 cm additionally refer to the importance of a certain successional stage for the species, as females of the species prefer low, shrubby oaks for oviposition (Ulrich and Caspari 2007).

Grazing can affect the habitat quality for insects by slowing down succession speed. Game grazing occurs primarily in palatable early and light successional stages (Treiber 2003; Joys et al. 2004; Hédl et al. 2010). Therefore, the influence of grazing animals after coppice abandonment may be crucial to maintaining light forest structures (Benes et al. 2006). In the coppice of our study area, regular Red Deer grazing keeps the oak trees in a bush-like stage, and therefore in a suitable condition for *S. ilicis* as larvae host plants (Köstler 2005). The oviposition occurs preferably on young and small oak trees, often less than 50 cm in height (Koschuh and Fauster 2005; Maes et al. 2014). However, conditions may change when intense grazing decreases the amount of fresh leaves and buds, which are the food source for caterpillars (Koschuh and Fauster 2005; Maes et al. 2014). Ulrich and Caspari (2007) observed that heavily grazed gnarled central shoots were even preferentially approached by *S. ilicis* for oviposition. This is in line with our results of increasing population density of *S. ilicis* with increasing Red Deer density. The present study confirms earlier studies concluding that *S. ilicis* benefits from a high density of grazers that maintain a certain habitat quality (Schiess-Bühler 2004; Hermann 2007; Ulrich and Caspari 2007; FVA-BW 2022).

For *M. athalia*, on the other hand, there was no detectable influence of grazing intensity on population density of the imagines. High Red Deer grazing pressure results in grass-rich stands (Gill and Beardall 2001; Kirby 2001; Gill and Fuller 2007) at the cost of dwarf shrubs such as *Calluna vulgaris* and Bilberry (*Vaccinium myrtillus*). Bilberry carpets often host larger populations of *M. athalia’s* larval food plant, *Melampyrum pratense* that is hemiparasite on Bilberry. *M. pratense* is not directly grazed but sensitive to trampling (Klotz and Kühn 2002). Intensive Red Deer grazing can also lead to damage of nectar plants and therefore can have a negative impact on butterflies (Feber 2001, Boulanger et al. 2018).

## Conclusions

*M. athalia* and *S. ilicis*, as light forest species, indicate the conservation value of young forest succession stages which provide habitats for many other insect species (Freese et al. 2006; Fartmann et al. 2013). Due to the decline of light-demanding insects in forests, the preservation of open forest systems with a mosaic of successional stages, such as coppice, is of particular importance (Freese et al. 2006; van Swaay et al. 2006; Benes et al. 2006; Vacik et al. 2009). Because of the preference for early successional stages in both species, a transformation of the studied coppice systems to spruce plantations or high beech forest would likely lead to a disappearance of both species in the area.

Conservation managers should aim to maintain coppice systems with large number of moderately isolated, large coupes of several hectares in size as in our study area, because the effects of fragmentation seem to be less important in such “intact” coppice systems. Habitat quality should be maintained through short rotation cycles that always provide a supply of well-connected, four-to six-year-old, large, continuous coupes, benefitting both species. For *S. ilicis*, a certain level of Red Deer grazing is beneficial. For *M. athalia*, very high Red Deer densities should be avoided. Climate change may reduce the habitat for *M. athalia* that meets its niche requirements, as it appears to be a more cold-adapted species in our area, suggesting that maintaining open forests with a high habitat quality is even more important.

## Acknowledgments

We are very grateful to the heads of the coppice management collectives of all villages in the coppice woods of the Lahn-Dill district, Hesse for allowing access to their area and providing a large amount of information, especially Johannes Eckhardt, Roger Weitzel, Dietmar Orth and Klaus-Dieter Schmidt. Harro Schäfer and Rolf Twardella provided valuable insight into historical and current coppice management and Lepidoptera communities of the study area. We thank the Hessische Landesamt für Naturschutz, Umwelt und Geologie and Landesbetrieb (HLNUG, HessenForst) and the Landesamt für Natur, Umwelt und Verbraucherschutz North Rhine-Westphalia (LANUV) for granting permits. Many thanks to Claudia Frank and Josef Kallmayer who gave statistical advice. J. Kamp thanks Norbert Hölzel for allowing him to work on this unfunded project while based at the University of Münster.

## Authors Contribution

A. Graser, M. Kelling, R. Pabst, M. Schulz and J. Kamp designed the original research. A. Graser, M. Kelling, R. Pabst and M. Schulz carried out fieldwork and prepared the draft of the article. A. Graser was responsible for writing up the article. M. Kelling and M. Schulz were mainly responsible for researching primary literature. A. Graser and R. Pabst did most of the analytic work. J. Kamp reviewed and supervised.

