## Supplemental Figure S1, S2 and Table S3, S4 for "Habitat quality, not patch isolation, drives distribution and abundance of two light-demanding butterflies in fragmented coppice landscapes"

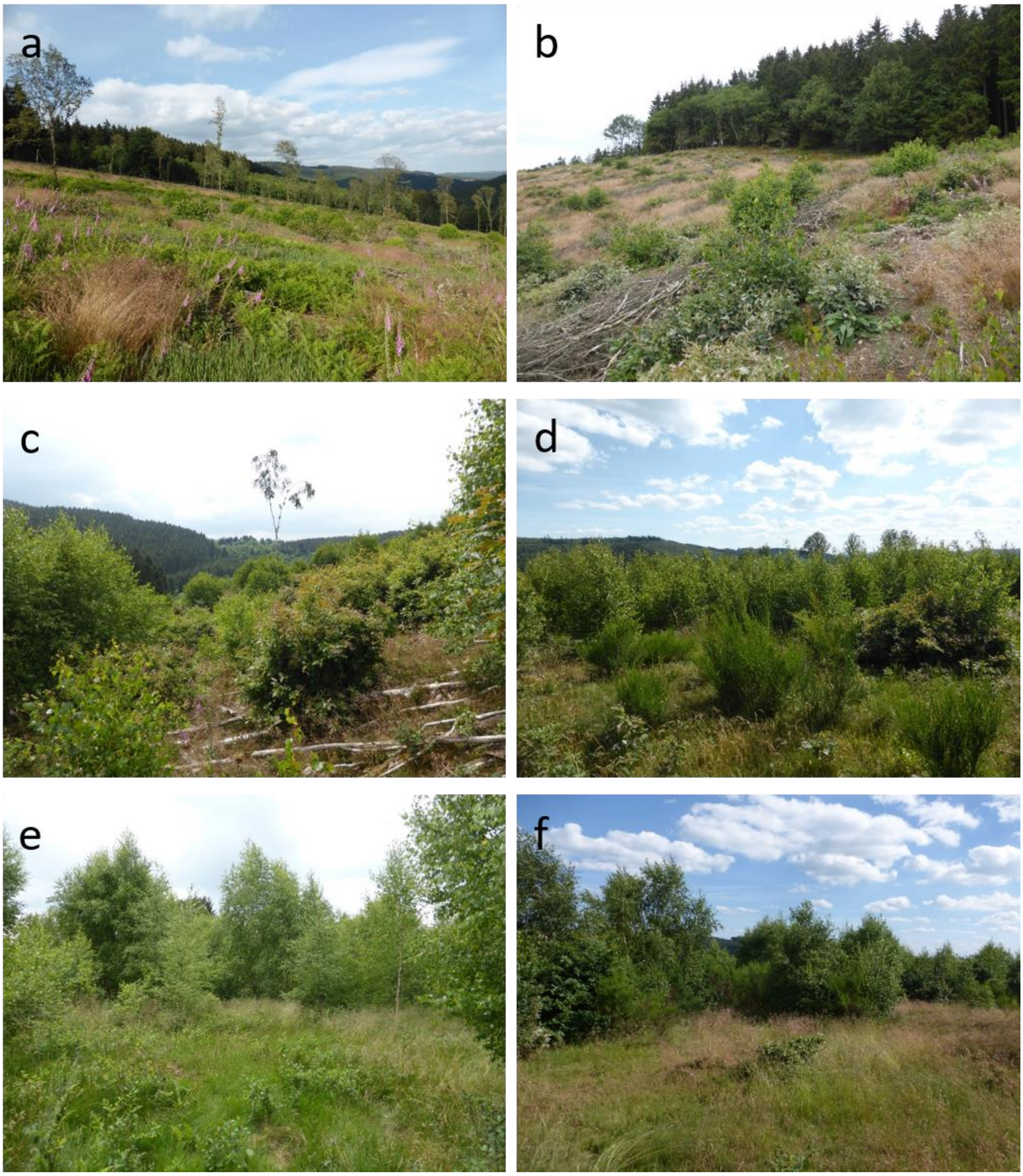


**Fig. S1**: Impressions of typical coppice stands: 1–2 years (a, b), 3–6 years (c, d), and 7–9 years after the last cut (e, f).

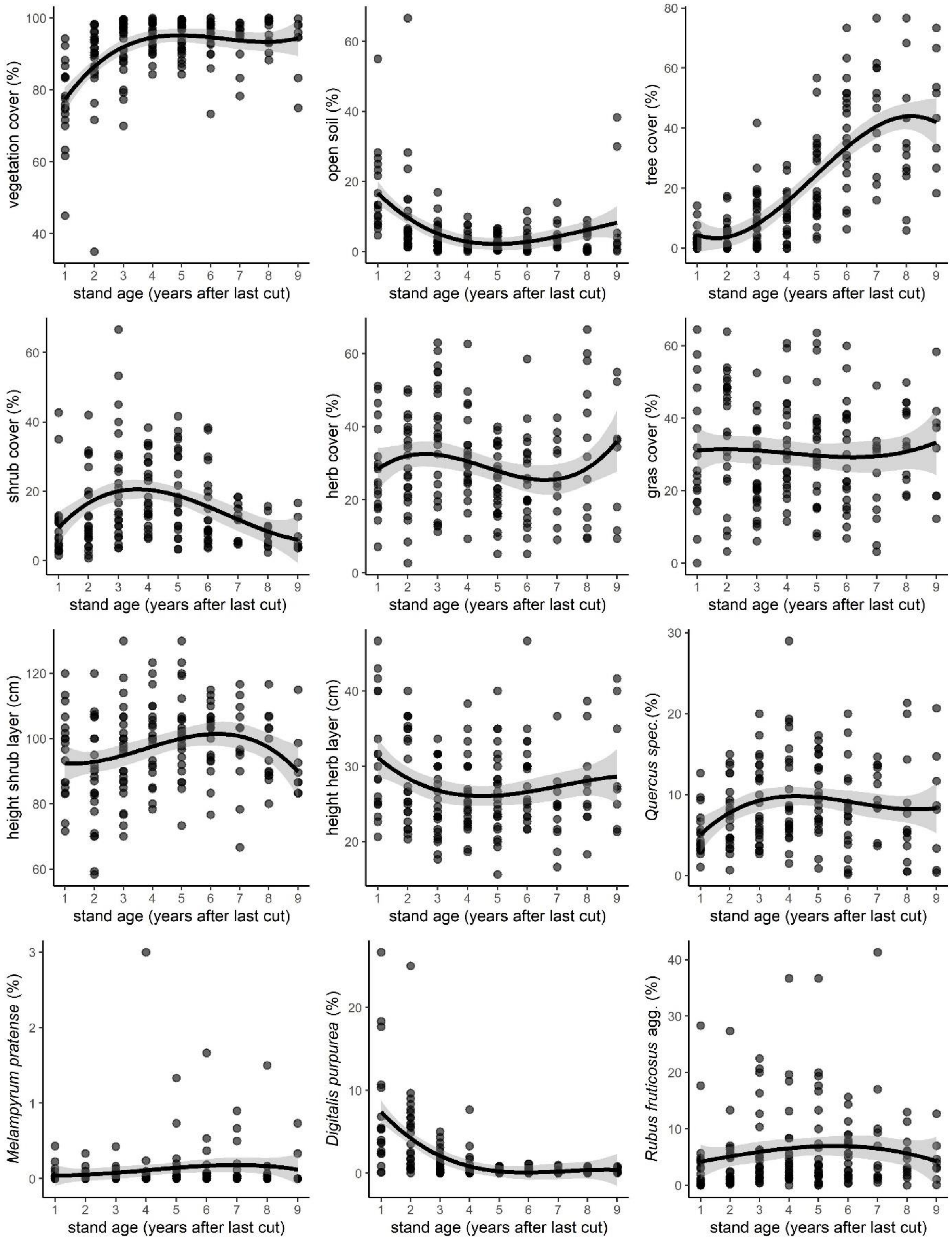


**Fig. S2**: Habitat conditions along successional gradient (1 – 9 years after last cut), scatterplot of raw data, the fitted line is a linear model with spline smoother.

**Table S3:** Habitat characteristics along the successional gradient of 1–9 years since last cut. All variables were predicted as a function of stand age in GLMMs, with the results illustrating significant associations of successional stage and measured habitat variables.

| **Model** | **Parameter** | **Distribution family** | **Estimate** | **SE** | **p** | | **AIC** |
|---|---|---|---|---|---|---|---|
| Total vegetation cover | Intercept | poisson | 1.43779 | 0.04601 | <0.0001 | *** | 711.8 |
| | Total vegetation (% cover) | | 0.28520 | 0.05386 | <0.0001 | *** | |
| Tree cover | Intercept | compois | 1.46084 | 0.05671 | <0.0001 | *** | 603.8 |
| | Trees (% cover) | | 0.54503 | 0.04230 | <0.0001 | *** | |
| | Trees ^2 (% cover) | | -0.13403 | 0.02540 | <0.0001 | *** | |
| Shrub cover | Intercept | poisson | 1.44869 | 0.04564 | <0.0001 | *** | 740.6 |
| | Shrubs (% cover) | | -0.07691 | 0.04415 | 0.0815 | . | |
| Herb cover | Intercept | poisson | 1.39598 | 0.05866 | <0.0001 | *** | 743.1 |
| | Herbs (% cover) | | -0.04631 | 0.04424 | 0.295 | n.s. | |
| | Herbs^2 (% cover) | | 0.05115 | 0.03417 | 0.134 | n.s. | |
| Gras cover | Intercept | poisson | 1.51251 | 0.05889 | <0.0001 | *** | 742.9 |
| | Gasses (% cover) | | 0.01446 | 0.04453 | 0.745 | n.s. | |
| | Grasses^2 (% cover) | | -0.06343 | 0.03886 | 0.103 | n.s. | |
| Deadwood cover | Intercept | genpois | 1.26849 | 0.05615 | <0.0001 | *** | 676.6 |
| | Deadwood (% cover) | | -0.35884 | 0.04348 | <0.0001 | *** | |
| | Deadwood^2 (% cover) | | 0.10619 | 0.02408 | <0.0001 | *** | |
| Open soil | Intercept | poisson | 1.40750 | 0.05113 | <0.0001 | *** | 728.4 |
| | Soil (% cover) | | -0.30799 | 0.08562 | 0.000322 | *** | |
| | Soil^2 (% cover) | | 0.03794 | 0.01952 | 0.051938 | . | |
| Height shrub | Intercept | poisson | 1.50581 | 0.05423 | <0.0001 | *** | 740.2 |
| | Shrub layer height (cm) | | 0.05898 | 0.04416 | 0.1817 | n.s. | |
| | Shrub layer height ^2 (cm) | | -0.05904 | 0.03262 | 0.0703 | . | |
| Height herb | Intercept | poisson | 1.44834 | 0.04600 | <0.0001 | *** | 742.4 |
| | Herb layer height (cm) | | -0.04818 | 0.04311 | 0.264 | n.s. | |
| *Quercus* spec. | Intercept | poisson | 1.44824 | 0.04597 | <0.0001 | *** | 741.7 |
| | *Quercus* spec. (% cover) | | 0.05918 | 0.04216 | 0.16 | n.s. | |
| *Melampyrum pratense* | Intercept | poisson | 1.47071 | 0.04694 | <0.0001 | *** | 742.2 |
| | *Melampyrum pratense* (% cover) | | 0.15854 | 0.08539 | 0.0634 | . | |
| | *Melampyrum pratense* ^2 (% cover) | | -0.01893 | 0.01364 | 0.1651 | n.s. | |
| *Digitalis purpurea* | Intercept | poisson | 1.29117 | 0.05211 | <0.0001 | *** | 665.3 |
| | *Digitalis purpurea* (% cover) | | -0.81722 | 0.10899 | <0.0001 | *** | |
| | *Digitalis purpurea* ^2 (% cover) | | 0.11387 | 0.02502 | <0.0001 | *** | |
| *Rubus fruticosus* agg | Intercept | poisson | 1.44844 | 0.04614 | <0.0001 | *** | 743.4 |
| | *Rubus fruticosus* agg (% cover) | | 0.02158 | 0.04272 | 0.613 | n.s. | |

**Table S4**: Correlations of the habitat parameters with PCA axis 1 and 2.

| **Habitat parameter** | **Axis** | |
|---|---|---|
| | **PC1** | **PC2** |
| Vegetation cover (%) | **0.99855795** | **-1.1864961** |
| Cover tree layer (%) | **1.34607218** | 0.26968031 |
| Cover shrub layer (%) | 0.44819407 | 0.22053061 |
| Cover herb layer (%) | **-0.8149916** | **-1.4241503** |
| Cover herbs (%)* | -0.2178492 | **-0.9296073** |
| Cover grasses (%)* | -0.7556437 | **-0.8048881** |
| Deadwood (%) | -0.6953617 | 0.79360701 |
| Open soil (%) | -0.7299732 | **1.22136971** |
| Height herb layer (cm) | -0.5864983 | -0.0586668 |
| Height shrub layer (cm) | 0.48056216 | 0.2416018 |
| *Melampyrum pratense (%)* | 0.35553749 | 0.02444958 |
| *Digitalis purpurea (%)* | **-0.8057455** | 0.64823911 |
| *Quercus spec (%)* | **0.93609155** | 0.18334194 |
| *Rumex acetosa (%)* | -0.3010535 | -0.1373987 |
| *Rubus rubus (%)* | 0.2957196 | 0.16634356 |
| *Galium harcynicum (%)* | 0.0058561 | -0.7748679 |
| *Betula spec (%)* | **1.4088236** | 0.20382503 |
| *Vaccinium myrtillus (%)* | -0.0954398 | -0.5858935 |
| *Calluna vulgaris (%)* | 0.48604324 | -0.0877497 |

*Proportion of the herb cover
